# A large-scale systematic survey of SARS-CoV-2 antibodies reveals recurring molecular features

**DOI:** 10.1101/2021.11.26.470157

**Authors:** Yiquan Wang, Meng Yuan, Jian Peng, Ian A. Wilson, Nicholas C. Wu

## Abstract

In the past two years, the global research in combating COVID-19 pandemic has led to isolation and characterization of numerous human antibodies to the SARS-CoV-2 spike. This enormous collection of antibodies provides an unprecedented opportunity to study the antibody response to a single antigen. From mining information derived from 88 research publications and 13 patents, we have assembled a dataset of ∼8,000 human antibodies to the SARS-CoV-2 spike from >200 donors. Analysis of antibody targeting of different domains of the spike protein reveals a number of common (public) responses to SARS-CoV-2, exemplified via recurring IGHV/IGK(L)V pairs, CDR H3 sequences, IGHD usage, and somatic hypermutation. We further present a proof-of-concept for prediction of antigen specificity using deep learning to differentiate sequences of antibodies to SARS-CoV-2 spike and to influenza hemagglutinin. Overall, this study not only provides an informative resource for antibody and vaccine research, but fundamentally advances our molecular understanding of public antibody responses to a viral pathogen.

## INTRODUCTION

From the beginning of COVID-19 pandemic, many research groups worldwide turned their attention to SARS-CoV-2 and, in particular, to the immune response to infection and vaccination. Over the past two years, thousands of human monoclonal antibodies to SARS-CoV-2 have been isolated and characterized [1, 2]. The major surface antigen to which antibodies are elicited is the SARS-CoV-2 spike (S) protein, which is a homotrimeric glycoprotein that facilitates virus entry by first engaging the host receptor ACE2 and then mediating membrane fusion [3, 4]. The S protein has three major domains, namely the N-terminal domain (NTD), receptor-binding domain (RBD), and S2 domain [5, 6]. Most studies on SARS-CoV-2 antibodies have focused on the immunodominant RBD [7], because neutralizing antibodies can be elicited to it with very high potency [8, 9]. Antibodies to the NTD and the highly conserved S2 domain have also been discovered, but usually exhibit lower neutralizing potency [10-16].

A common or public antibody response describes antibodies to the same antigen in different donors that share genetic elements that usually result in similar modes of antigen recognition. Deciphering public responses to particular antigens is not only critical for uncovering the molecular features of recurring antibodies within the diverse antibody repertoire at the population level, but also important for development of effective vaccines [17, 18]. A conventional approach to study public antibody responses is to identify public clonotypes, which are antibodies from different donors that share the same immunoglobulin heavy variable (IGHV) gene and with similar complementarity-determining region (CDR) H3 sequences [19-23]. While this definition of public clonotypes has improved our understanding of public antibody response, it generally ignores the contribution of the light chain. Moreover, our recent study has shown that a public antibody response to influenza hemagglutinin is driven by an IGHD gene with minimal dependence on the IGHV gene [24]. Therefore, the true extent and molecular characterization of public antibody responses remain to be explored.

Although information of many human clonal antibodies to SARS-CoV-2 is now publicly available, it has been difficult to leverage all available information to investigate public antibody responses to SARS-CoV-2. One major challenge is that the data from different studies are rarely in the same format. This inconsistency imposes a huge barrier to data mining. The establishment of the coronavirus antibody database (CoV-AbDab) has enabled researchers to deposit their antibody data in a standardized format and has partially resolved the data formatting issue [2]. However, not every SARS-CoV-2 antibody study has deposited their data to CoV-AbDab. Furthermore, IGHD gene identities, nucleotide sequences, and donor IDs are not available in CoV-AbDab, which makes it challenging to study public antibody responses using CoV-AbDab. Thus, additional efforts must be made to fully synergize the information across many different SARS-CoV-2 antibody studies to investigate and decipher public antibody responses.

In this study, we performed a systematic literature survey and assembled a large dataset of human SARS-CoV-2 monoclonal antibodies with donor information. We then analyzed this dataset and uncovered many previously unknown antibody sequence features that contribute to public antibody responses to SARS-CoV-2 S. For example, we identified a public antibody response to RBD that is largely independent of the IGHV gene, as well as involvement of a particular IGHD gene in a public antibody response to S2. Our analysis also revealed a number of recurring somatic hypermutations (SHMs) in different public clonotypes.

## RESULTS

### Collection of SARS-CoV-2 antibody information

Information for 8,048 human antibodies was collected from 88 research publications and 13 patents that described the discovery and characterization of antibodies to SARS-CoV-2 (**Figure S1, Data S1**). Among these antibodies, which were isolated from 215 different donors, 7,997 (99.4%) react with SARS-CoV-2, and the remaining 51 react with SARS-CoV or seasonal coronaviruses. While 99.1% (7,923/7,997) SARS-CoV-2 antibodies in our dataset bind to S protein, 49 bind to N and 25 to ORF8. Epitope information was available for most SARS-CoV-2 S antibodies, with 5,002 to RBD, 513 to NTD, and 890 to S2. In addition, information on neutralization activity, germline gene usage, sequence, structure, bait for isolation (e.g. RBD, S), and donor status (e.g. infected patient, vaccinee, etc.), if available, was collected for individual antibodies.

### Epitope-dependent V gene usage bias in SARS-CoV-2 S antibodies

To identify the sequence features in RBD, NTD, and S2 antibodies, we first performed an analysis on V gene usage. Our analysis identified several commonly used IGHV/IGK(L)V pairs among RBD antibodies (**Figure 1A**), such as IGHV3-53/IGKV1-9 and IGHV3-53/IGKV3-20, which represent two known public clonotypes [25-30]. We also observed substantial enrichment of IGHV1-24 among NTD antibodies over the naïve baseline (**Figure 1B**), which was established by published datasets of antibody repertoire sequencing from 26 healthy donors [31-33]. IGHV1-24 is in fact a known public antibody response that targets an antigenic supersite on NTD [10-13]. These observations illustrate that the gene usage pattern in our dataset is consistent with previous findings. Importantly, our dataset also enabled us to discover previously unknown patterns in gene usage. For example, IGHV3-30 and IGHV3-30-3 were highly enriched among S2 antibodies over baseline (**Figure 1B**). For our subsequent analyses, IGHV3-30-3 was also labeled as IGHV3-30, since IGHV3-30 and IGHV3-30-3 have an identical amino acid sequence in the framework regions, CDR H1 and CDR H2. V gene usage bias was also observed in the light chain. For example, IGKV3-20 and IGKV3-11 were most used among S2 antibodies, whereas IGKV1-33 and IGKV1-39 were most used among RBD antibodies (**Figure 1C**). Overall, these results demonstrated that RBD, NTD, and S2 antibodies have distinct patterns of V gene usage.

**Figure 1.**
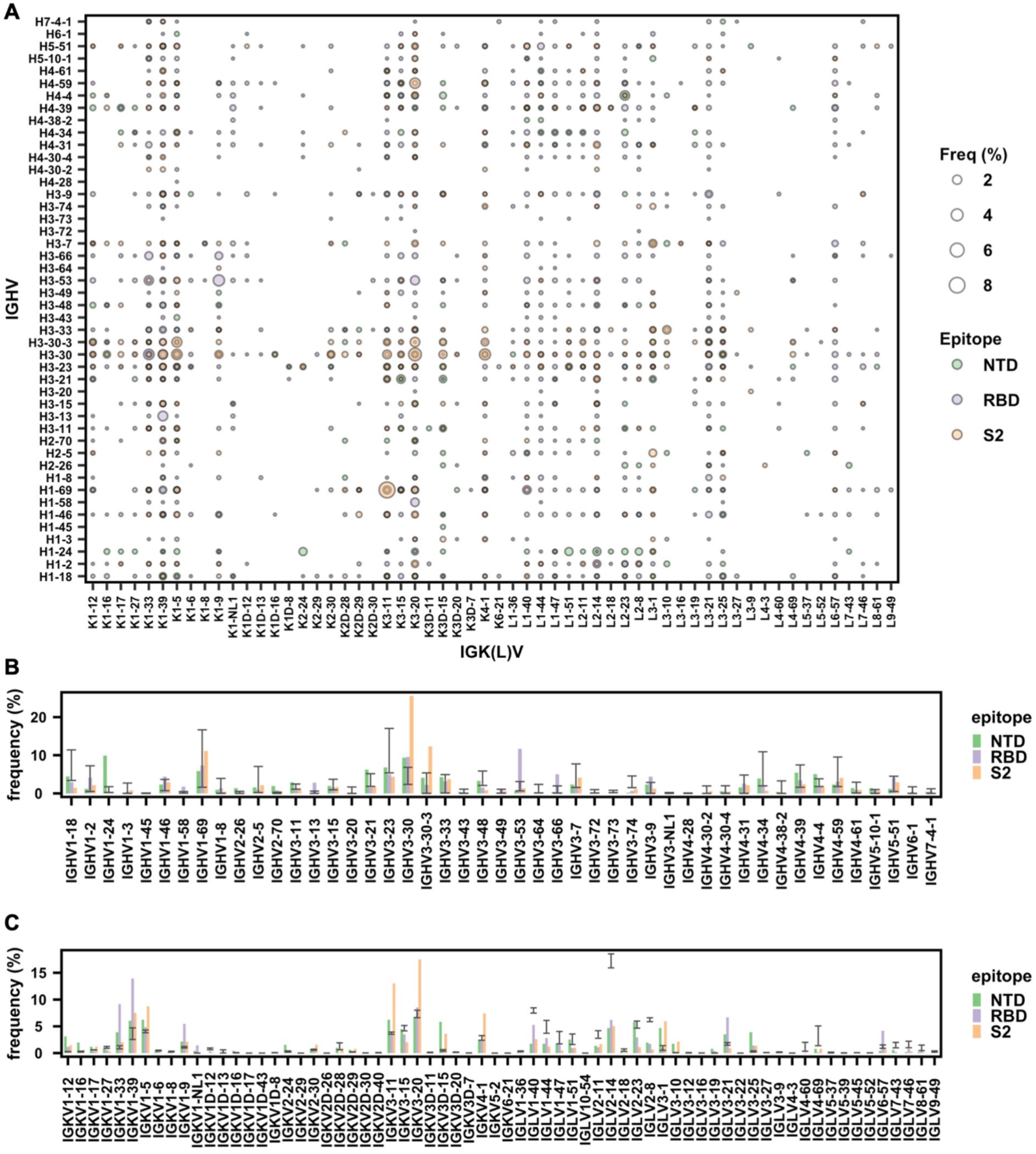
Analysis of V gene usage in SARS-CoV-2 S antibodies. **(A)** The frequency of different V gene pairings between heavy and light chains are shown for SARS-CoV-2 S antibodies to RBD, NTD, and S2. The size of each datapoint represents the frequency of the corresponding IGHV/IGK(L)V pair within its epitope category. Only those antibodies where both IGHV and IGK(L)V information is available for both heavy and light chains was included in this analysis. **(B)** The IGHV gene usage in antibodies to NTD, RBD, and S2 are shown. Only those antibodies with IGHV information available were included in this analysis. **(C)** The IGK(L)V gene usage in antibodies to NTD, RBD, and S2 are shown. Only those antibodies with IGK(L)V information available were included in this analysis. **(B-C)** Error bars represent the frequency range among 26 healthy donors [31-33].

### CDR H3 analysis reveals public antibody response

Although heavy and light chain V genes together encode four of the six CDRs, most of the antibody sequence diversity comes from the CDR H3 region due to V(D)J recombination. Since CDR H3 is typically an important determinant for binding and may even dominate the paratope [24, 34-37], characterization of CDR H3 sequences in S antibodies is essential for understanding the antibody response to SARS-CoV-2. Here, we aimed to examine the convergence of CDR H3 sequences among S antibodies. Briefly, CDR H3 sequences with the same length were clustered by an 80% sequence identity cutoff. Only those clusters that contained antibodies from at least two different donors were subjected to further analysis. A total of 170 clusters were identified (**Figure 2A and Data S1**). Interestingly, antibodies within the same cluster often share the same binding region on the S protein (RBD, NTD, or S2), consistent with the notion that the CDR H3 sequence has a critical role in determining the epitope that is recognized.

**Figure 2.**
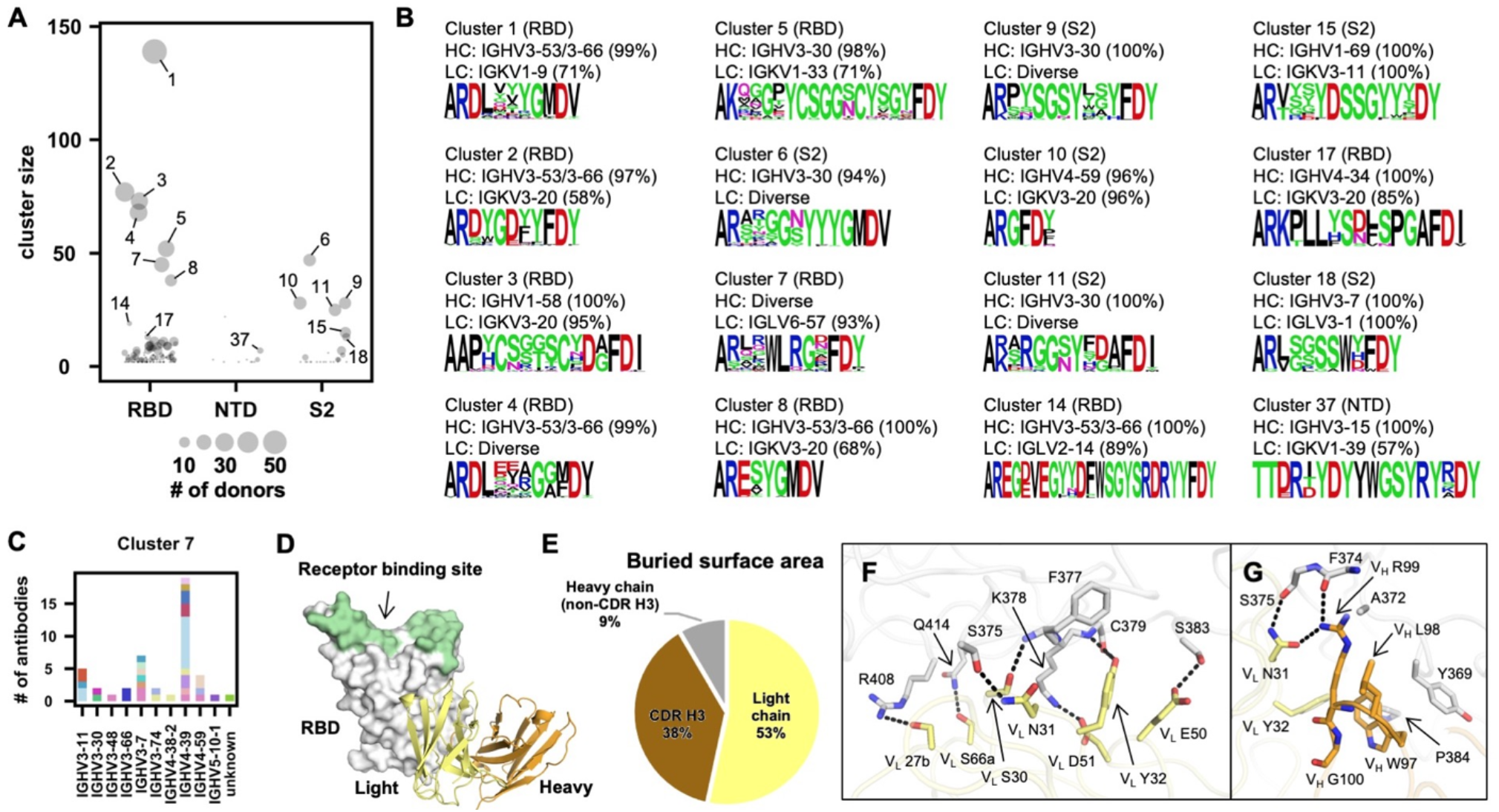
Convergent CDR H3 sequences among SARS-CoV-2 S antibodies. **(A)** CDR H3 sequences from individual antibodies were clustered using a 20% cutoff (see Materials and Methods). The epitope of each CDR H3 cluster is classified based on that of its antibody members. Cluster size represents the number of antibodies within the cluster. **(B)** The V gene usage and CDR H3 sequence are shown for each of the 16 CDR H3 clusters of interest. For each of the CDR H3 cluster of interest, the CDR H3 sequences are shown as a sequence logo, where the height of each letter represents the frequency of the corresponding amino-acid variant (single-letter amino-acid code) at the indicated position. The dominant germline V genes (>50% usage among all antibodies within a given CDR H3 cluster) are listed. Diverse: no germline V genes had >50% frequency among all antibodies within a given CDR H3 cluster. HC: heavy chain. LC: light chain. **(C)** IGHV usage in cluster 7 is shown. Different colors represent different donors. Unknown: IGHV information is not available. **(D)** An overall view of SARS-CoV-2 RBD in complex with IGLV6-57 antibody S2A4 (PDB 7JVA) [41], which belongs to cluster 7, is shown. The RBD is in white with the receptor binding site highlighted in green. The heavy and light chains of S2A4 are in orange and yellow, respectively. **(E)** Percentages of the S2A4 epitope that are buried by the light chain, heavy chain (without CDR H3), and CDR H3 are shown as a pie chart. Buried surface area (BSA) was calculated by PISA (Proteins, Interfaces, Structures and Assemblies) at the European Bioinformatics Institute (https://www.ebi.ac.uk/pdbe/prot_int/pistart.html) [74]. **(F-G)** Detailed interactions between the **(F)** light and **(G)** heavy chains of S2A4 and SARS-CoV-2 RBD. Hydrogen bonds and salt bridges are represented by black dashed lines. The color coding is the same as panel D.

The largest cluster (cluster 1) consisted of 139 antibodies from 57 donors (**Figure 2B**). Most of the antibodies in cluster 1 belonged to a well-characterized public clonotype to RBD that is encoded by IGHV3-53/3-66 and IGKV1-9 [25-27, 29, 30]. IGHV3-53/3-66, which is frequently used in RBD antibodies [28], was also enriched among antibodies in several other major CDR H3 clusters (e.g. clusters 2, 4, 8, and 14). Antibodies that bind to quaternary epitopes by bridging two RBDs on the same spike are found in clusters 14 and 17 [38] (**Figure S2**). Notably, both clusters 3 and 5, which target the RBD, contained a conserved disulfide bond (**Figure 2B**). Cluster 3 represents another well-characterized public clonotype that is encoded by IGHV1-58/IGKV3-20 [8, 9, 39, 40]. On the other hand, antibodies in cluster 5, which are largely encoded by IGHV3-30/IGKV1-33, have not been extensively studied. Most antibodies within cluster 5 had relatively weak neutralizing activity, if any, despite having reasonable binding affinity (**Table S1**). This result suggests the existence of an RBD-targeting public clonotype that had minimal neutralizing activity. Similar observation was made with RBD antibodies encoded by IGHV3-13/IGKV1-39, although most of these antibodies did not share a similar CDR H3 (**Figure S3 and Table S2**).

Furthermore, we also discovered several S2-specific CDR H3 clusters (clusters 6, 9, and 11) that were predominantly encoded by IGHV3-30 with diverse IGK(L)V genes, suggesting a public heavy chain response to S2 (**Figure 2B**). Clusters 10 and 15 were also of interest to us. Cluster 10 was featured by a very short CDR H3 (6 amino acids, IMGT numbering) and was encoded by IGHV4-59/IGKV3-20, which was a frequent V gene pair among the S2 antibodies. Cluster 15 was encoded by IGHV1-69/IGKV3-11, which was the most used V gene pair among the S2 antibodies. Therefore, clusters 10 and 15 represented two major S2 public clonotypes, despite their minimal neutralizing activity (**Table S1**). In contrast to RBD- and S2-specific clusters, all NTD-specific CDR H3 clusters had a relatively small size (**Figure 2A**), suggesting that the paratopes for most NTD antibodies are not dominated by CDR H3.

### A public antibody response dominated by the light chain and CDR H3

While most clusters have a dominant IGHV gene, diverse IGHV genes were observed in cluster 7 (**Figure 2B-C**). Most antibodies (42 out of 45) in cluster 7 used IGLV6-57, suggesting their paratopes are mainly composed of CDR H3 and light chain. S2A4, which is encoded by IGHV3-7/IGLV6-57 [41], is an antibody in cluster 7. A previously determined structure of S2A4 in complex with RBD indeed demonstrates that its CDR H3 contributes 38% of the buried surface area (BSA) of the epitope, whereas the light chain contributes 53% (**Figure 2D-E**). Specifically, IGLV6-57 forms an extensive H-bond network with the RBD (**Figure 2F**), whereas a ^97^WLRG^100^ motif at the tip of CDR H3 interacts with the RBD through H-bonds, π-π stacking, and hydrophobic interactions (**Figure 2G**). Although G100 does not participate in binding, it exhibits backbone torsion angles (Φ = −94°, Ψ = −160°) that are in the preferred region of Ramachandran plot for glycine, but in the allowed region for non-glycine (**Figure S4**). Consistently, this ^97^WLRG^100^ motif is highly conserved in cluster 7 (**Figure 2B**). These results illustrate that our CDR H3 clustering analysis not only captured existing knowledge about public SARS-CoV-2 antibody responses, but was able to uncover recurring sequence features among SARS-CoV-2 antibodies that were previously unknown.

### IGHV3-30/IGHD1-26 is a recurring feature in S2 antibodies

As a major contributor to CDR H3, the IGHD gene can also drive a public antibody response [24]. Consequently, we aimed to understand if there are any signature IGHD genes in SARS-CoV-2 S antibodies. While the frequency of most IGHD genes were within the baseline level, IGHD1-26 was highly enriched among S2 antibodies (**Figure 3A**). These IGHD1-26 S2 antibodies were predominantly encoded by IGHV3-30 (**Figure 3B**), which is one of the most used IGHV genes among S2 antibodies (**Figure 1B**). In contrast, the IGK(L)V gene usage was more diverse among these IGHD1-26 S2 antibodies, although several were more frequently used than others (**Figure 3C**), implying that this public antibody response to S2 is mainly driven by the heavy chain. Interestingly, 70% of these IGHD1-26 S2 antibodies had a CDR H3 of 14 amino acids, whereas only <20% of other S antibodies had a CDRH3 of 14 amino acids (**Figure 3D**). In fact, most members of clusters 6, 9, and 11 in our CDR H3 analysis above (**Figure 2B**) represented this public antibody response to S2. While CDR H3 is also encoded by the IGHJ gene, the distribution of IGHJ gene usage in these IGHD1-26 S2 antibodies did not show a strong deviation from that of other S antibodies in our dataset (**Figure 3E**).

**Figure 3.**
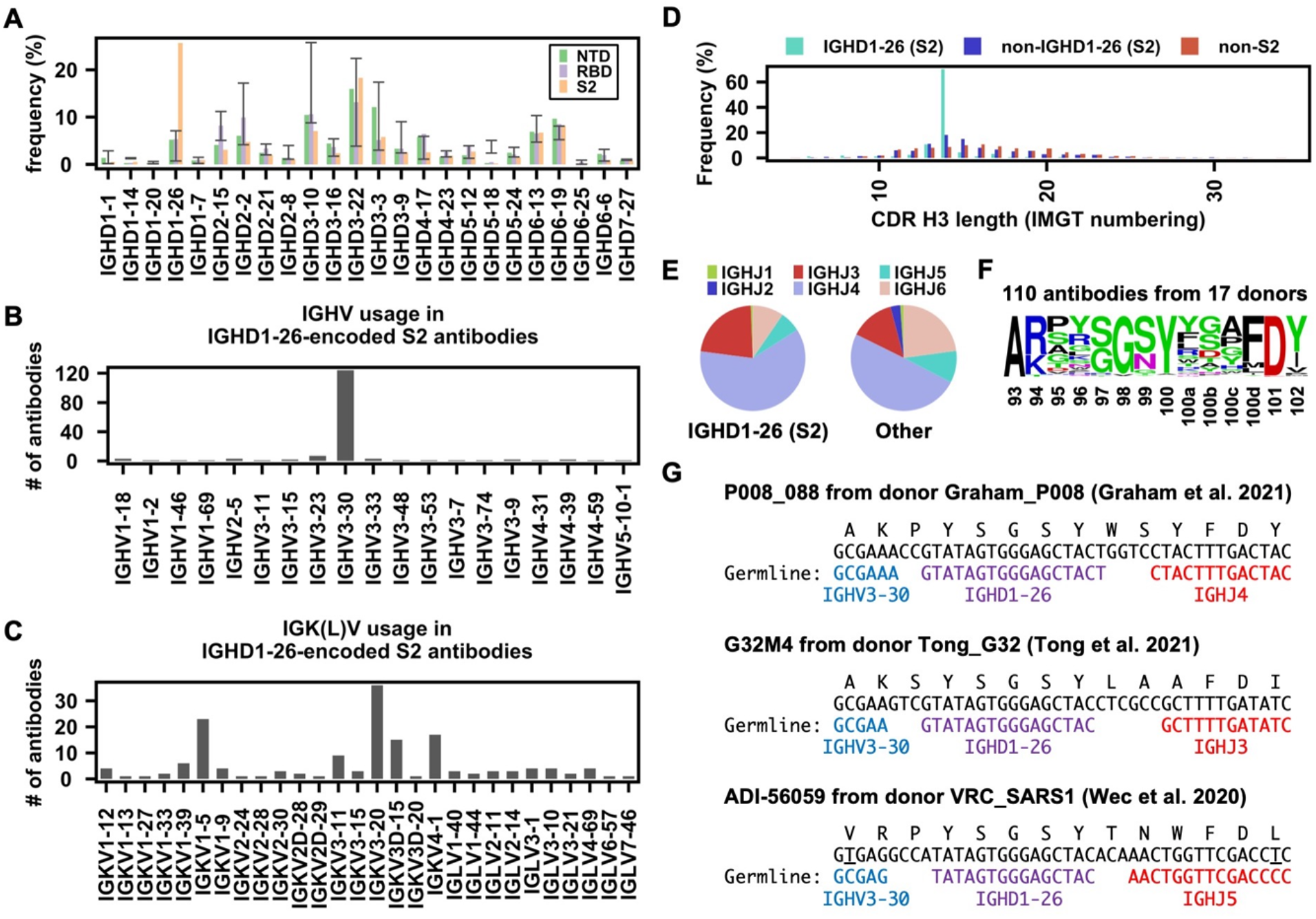
Enrichment of IGHD1-26 in SARS-CoV-2 S2 antibodies. **(A)** The IGHD gene usage in NTD, RBD, S2 antibodies is shown. Error bars represent the frequency range among 26 healthy donors. **(B)** IGHV gene usage and **(C)** IGK(L)V gene usage among IGHD1-26 S2 antibodies is shown (n = 157). **(D)** The distribution of CDR H3 length (IMGT numbering) in IGHD1-26 S2 antibodies (n = 157), non-IGHD1-26 S2 antibodies (n = 533), and other non-S2 S antibodies that do not target S2 (n = 5,090), are shown. **(E)** The IGHJ gene usage among IGHD1-26 S2 antibodies (n = 157) and other S antibodies with well-defined epitopes (n = 5,623) is shown. **(F)** The CDR H3 sequences for IGHD1-26 S2 antibodies (n = 110) are shown as a sequence logo. **(G)** Amino acid and nucleotide sequences of the V-D-J junction are shown for three IGHD1-26 S2 antibodies [42-44]. Putative germline sequences and segments were identified by IgBlast [66] and are indicated. Somatically mutated nucleotides are underlined. Intervening spaces at the V-D and D-J junctions are N-nucleotide additions.

In our dataset, there were 110 IGHD1-26 S2 antibodies from 17 donors with a CDR H3 length of 14 amino acids. Sequence logo analysis of these 110 antibodies revealed a conserved ^97^[S/G]G[S/N]Y^100^ motif in the middle of their CDR H3 sequences (**Figure 3F**). In-depth analysis of the CDR H3 sequences from three representative IGHD1-26 S2 antibodies, namely P008_088, G32M4, and ADI-56059, further indicated that the conserved ^97^[S/G]G[S/N]Y^100^ motif was within the IGHD1-26-encoded region (**Figure 3G**). Of note, P008_088, G32M4, and ADI-56059 were isolated from three different donors by three independent research groups [42-44]. While P008_088 and G32M4 were from SARS-CoV-2 infected individuals, ADI-56059 was from a SARS-CoV survivor. Although 87 out of these 110 IGHD1-26 S2 antibodies can cross-react with SARS-CoV, they generally have minimal neutralization activity (**Table S3**). Together, these results show that IGHV3-30/IGHD1-26 represents a public antibody response to a highly conserved epitope in S2.

### Recurring somatic hypermutations in public antibody responses

Our recent study has shown that V_H_ Y58F is a recurring somatic hypermutation (SHM) among IGHV3-53 antibodies to SARS-CoV-2 RBD [25]. Here, we aimed to identify additional recurring SHMs in other public clonotypes to SARS-CoV-2 S. In this analysis, antibodies from at least two donors that had the same IGHV/IGK(L)V genes and CDR H3s from the same CDR H3 cluster were classified as a public clonotype (**Figure 4A**). SHM that occurred in at least two donors within a public clonotypes was defined as a recurring SHM. Our analysis here only focused on major public clonotypes with antibodies from at least nine donors. This analysis led to the identification of several recurring SHMs in IGHV3-53/3-66-encoded public clonotypes that were previously characterized, including V_H_ F27V, T28I, and Y58F [25, 45, 46] (**Figure S5**). We also identified many other previously unknown recurring SHMs in both heavy and light chains (**Figure 4A-B**), including V_L_ S29R in a IGHV1-58/IGKV3-20 public clonotype that belongs to cluster 3 of our CDR H3 clustering analysis (**Figure 2A-B**). V_L_ S29R emerged in 8 out of 26 (31%) donors that carried this IGHV1-58/IGKV3-20 public clonotype.

**Figure 4.**
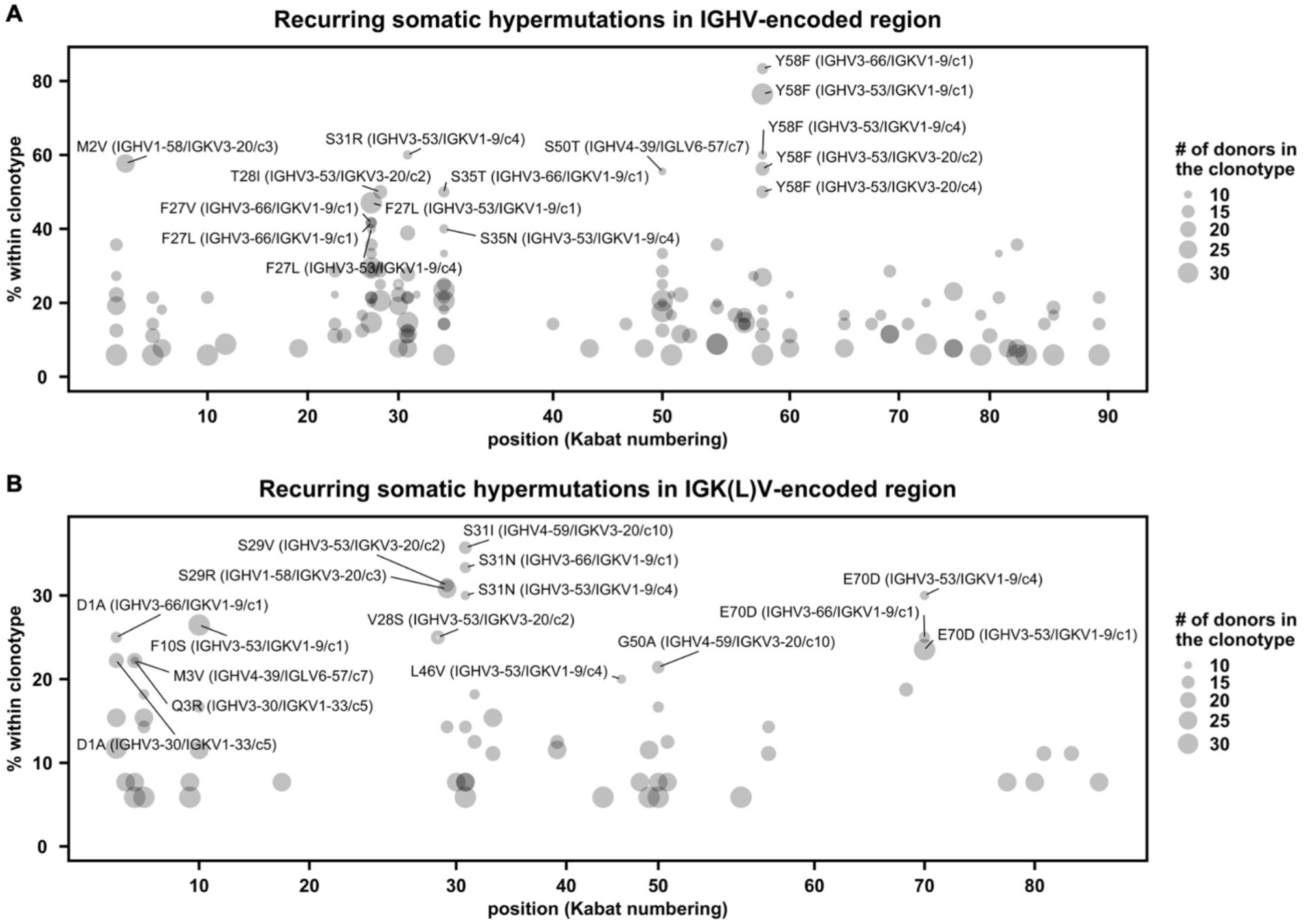
Recurring somatic hypermutations (SHMs) in SARS-CoV-2 S antibodies. **(A-B)** For each public clonotype, if the exact same SHM emerged in at least two donors, such SHM is classified as a recurring SHM. Only those public clonotypes that can be observed in at least nine donors are shown. **(A)** Recurring SHMs in heavy chain V genes. **(B)** Recurring SHMs in light chain V genes. X-axis represents the position on the V gene (Kabat numbering). Y-axis represents the percentage of donors who carry a given recurring SHM among those who carry the public clonotype of interest. For example, V_L_ S29R emerged in 8 donors out of 26 donors that carry an public clonotype that is encoded by IGHV1-58/IGKV3-20. As a result, V_L_ S29R (IGHV1-58/IGKV3-20) is 31% (8/26) within the corresponding clonotype. Of note, since each public clonotype is also defined by the similarity of CDR H3 (see Materials and Methods), there could be multiple clonotypes with the same heavy and light chain V genes (e.g. IGHV3-53/IGKV1-9). The CDR H3 cluster ID for each clonotype is indicated with a prefix “c”, following the information of the V genes. For heavy chain, SHMs that emerged in at least 40% of the donors of the corresponding clonotype are labeled. For light chain, SHMs that emerged in at least 20% of the donors of the corresponding clonotype are labeled.

Antibodies of this IGHV1-58/IGKV3-20 public clonotype bind to the ridge region of SARS-CoV-2 RBD (**Figure 5A**), and can be robustly elicited by infection with antigenically distinct variants of SARS-CoV-2 [39, 47] and by vaccination [48, 49]. These antibodies are also able to potently neutralize multiple variants of concern (VOC) [9, 48, 50]. We compared two previously determined structures of IGHV1-58/IGKV3-20 antibodies in complex with RBD [40, 51], where one has the germline-encoded V_L_ S29 (**Figure 5B**) and the other carries a somatically mutated V_L_ R29 (**Figure 5C**). While neither V_L_ S29 nor V_L_ R29 directly interact with RBD, V_L_ R29 is able to form a cation-π interaction with V_L_ Y32, which in turn forms a T-shaped π-π stacking with RBD-F486 and H-bonds with RBD-C480 (**Figure 5C**). In the absence of SHM V_L_ S29R, the rotamer adopted by V_L_ Y32 does not permit these interactions to be formed. During our structural analysis, we discovered that V_L_ S29R forms a salt bridge with another SHM V_L_ G92D (**Figure 5C**), which can further stabilize the interactions between V_L_ Y32 and with RBD. In fact, it is likely that V_L_ S29R promoted the emergence of V_L_ G92D, since V_L_ G92D was found in four out of the 67 antibodies and all four that carried V_L_ S29R (**Figure 5D-E**). This analysis substantiates the notion that recurring SHM can be found among antibodies within a public clonotype and further suggests the existence of common affinity maturation pathways that involve emergence of multiple SHMs in a defined order.

**Figure 5.**
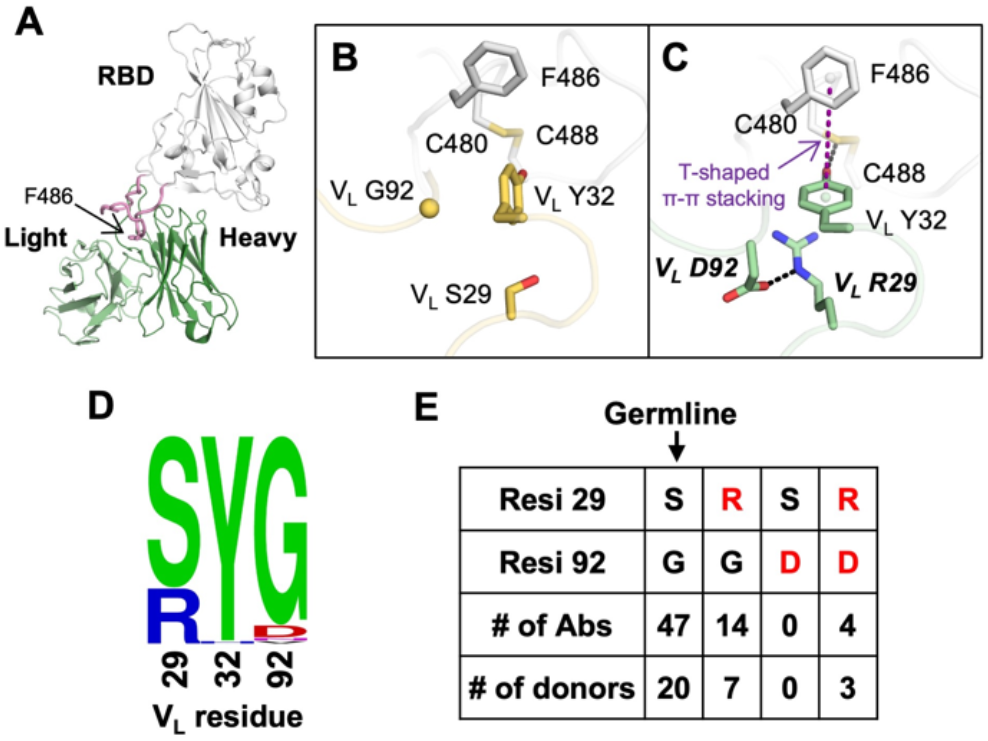
Structural analysis of a recurring SHM in the IGHV1-58/IGKV3-20 public clonotype. **(A)** An overall view of SARS-CoV-2 RBD in complex with the IGHV1-58/IGKV3-20 antibody PDI 222 (PDB 7RR0) [51]. The RBD is shown in white, while the heavy and light chains of the antibody are in dark and light green, respectively. The ridge region (residues 471-491) is shown in pink, with F486 highlighted as sticks. **(B-C)** Structural comparison between two IGHV1-58/IGKV3-20 antibodies that either **(B)** carry germline residues V_L_ S29/G92 (COVOX-253, PDB 7BEN) [40] and **(C)** somatically hypermutated residues V_L_ R29/D92 (PDI 222, PDB 7RR0) [51]. SARS-CoV-2 RBD is in white, while antibodies are in yellow (COVOX-253) and green (PDI 222). Somatically mutated residues are labeled with bold and italic letters. The T-shaped π-π stacking between RBD-F486 and V_L_ Y32 is indicated by a purple dashed line. Hydrogen bond and salt bridge are represented by black dashed lines. **(D)** Sequence logo of V_L_ residues 29, 32, and 92 among 67 IGHV1-58/IGKV3-20 RBD antibodies are shown. **(E)** Numbers of antibodies in the IGHV1-58/IGKV3-20 public clonotype carrying the germline-encoded variant at V_L_ residues 29 and 92 (S29, G92), as well as V_L_ SHM S29R and G92D (red) are listed. Of note, one antibody in this IGHV1-58/IGKV3-20 public clonotype carries S29/N92 and another carries S29/V92. However, they are not listed in the table here.

### Antigen identification by deep learning

Since many sequence features of public antibody responses to the S protein can be observed in our dataset, we postulated that the dataset is sufficiently large to train a deep learning model to identify S antibodies. To provide a proof-of-concept, we aimed to train a deep learning model to distinguish between antibodies to S and to influenza hemagglutinin (HA). Among different antigens, HA was chosen here because there are a large number of HA antibodies with published sequences, albeit still lower than the published SARS-CoV-2 S antibodies. Here, 4,736 unique SARS-CoV-2 S antibodies and 2,204 unique influenza HA antibodies with complete information for all six CDR sequences were used (**Data S2**). Sequences for HA antibodies were retrieved from GenBank [52]. None of these antibodies have identical sequences in all six CDRs. These antibodies to S and HA were divided into a training set (64%), a validation set (16%), and a test set (20%), with no overlap between the three sets. The training set was used to train the deep learning model. The validation set was used to evaluate the model performance during training. The test set was used to evaluate the performance of the final model.

Our deep learning model has a simple architecture, which consisted of one encoder per CDR followed by three fully connected layers (**Figure 6A**). To evaluate the model performance on the test set, the area under the curves of receiver operating characteristic (ROC AUC) and precision-recall (PR AUC) were used to measure the model’s ability to avoid misclassification. While ROC AUC is popular evaluation metric [53], PR AUC is shown to be more informative for evaluating models that are trained with imbalanced datasets [54]. Model performance was the best when all six CDRs (i.e. H1, H2, H3, L1, L2, and L3) were used to train the model, which resulted in an ROC AUC and an PR AUC of 0.87 and 0.92, respectively (**Figure 6B and Table S4**). Interesting, a similar performance was observed when the model was trained by the three heavy-chain CDRs (i.e. H1, H2, and H3) (ROC AUC = 0.86, PR AUC = 0.91), indicating that the heavy chain sequence captures most of the information to distinguish between HA antibodies and S antibodies. A reasonable performance was also observed when the model was trained by the three light-chain CDRs (i.e. L1, L2, and L3) (ROC AUC = 0.77, PR AUC = 0.86). For other types of inputs that we have tested, including CDR H3 only, CDR L3 only, CDR H3+L3, CDR H1+H2, and CDR L1+L2, the ROC AUCs were between 0.72 and 0.83 and the PR AUCs were between 0.82 and 0.90. These results imply that IGHV-encoded region (H1+H2), IGK(L)V-encoded region (L1+L2), and the V(D)J junctions (CDR H3 and CDR L3) are all informative for predicting antigen specificity. Overall, while our deep learning model had a relatively simple architecture, it was able to discriminate between antibodies to two different antigens based on primary sequences.

**Figure 6.**
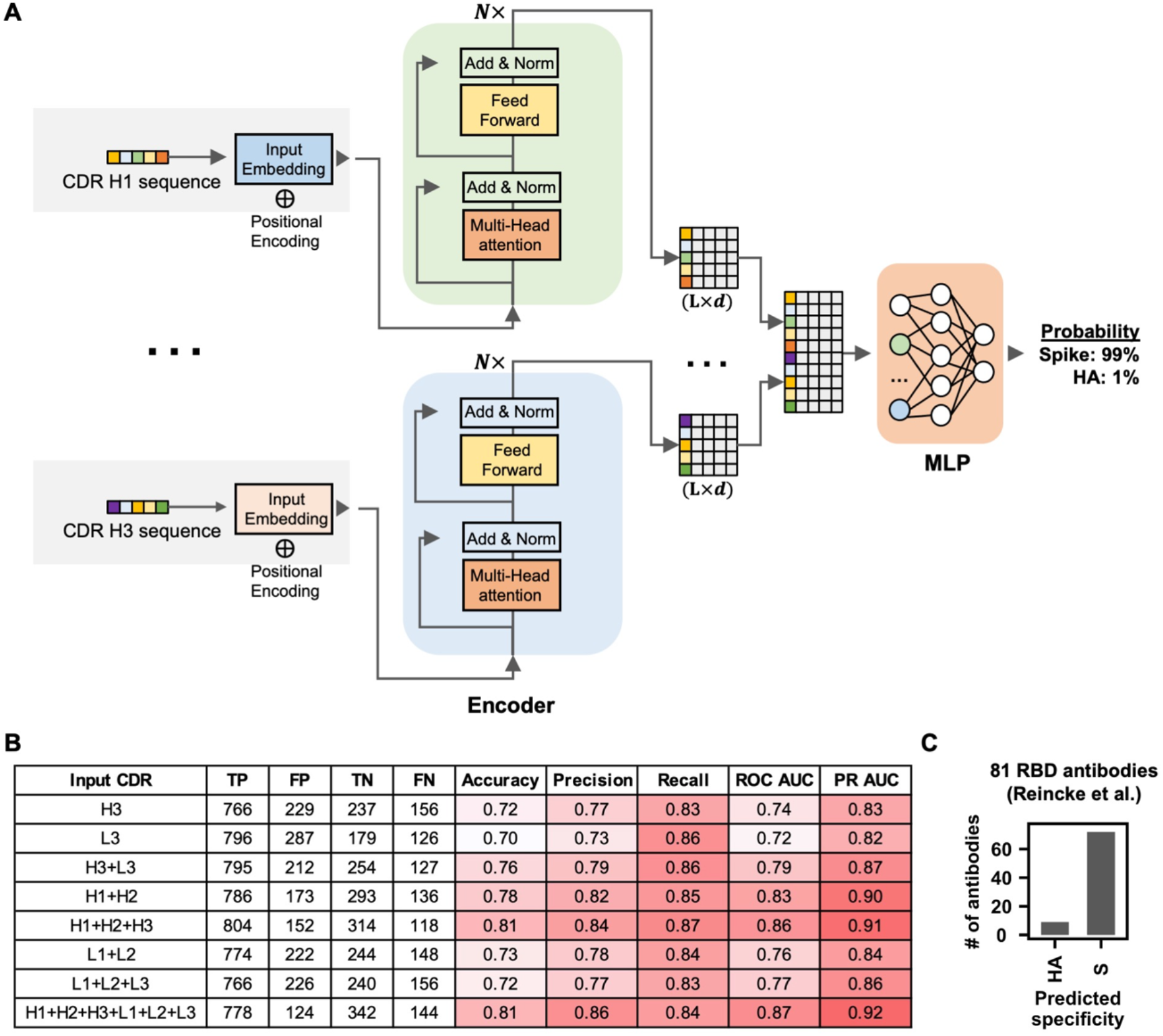
Antigen identification by deep learning. **(A)** A schematic overview of the deep learning model architecture. **(B)** For evaluating model performance, S antibodies and HA antibodies were considered “positive” and “negative”, respectively. Model performance on the test set was compared when different input types were used. Of note, the test set has no overlap with the training set and the validation set, both of which were used to construct the deep learning model. True positive (TP) represents the number of S antibodies being correctly classified as S antibodies. False positive (FP) represents the number of HA antibodies being misclassified as S antibodies. True negative (TN) represents the number of HA antibodies being correctly classified as HA antibodies. False negative (FN) represents the number of S antibodies being misclassified as HA antibodies. See Materials and Methods for the calculations of accuracy, precision, recall, ROC AUC, and PR AUC for the training and test sets. **(C)** The antigen specificity of 81 RBD antibodies from Reincke et al. [47] were predicted by a deep learning model that was trained to distinguish between S antibodies and HA antibodies.

A recent study reported 81 antibodies to SARS-CoV-2 RBD that were elicited by Beta variant infection [47]. While these 81 antibodies were not included in the dataset that we assembled (**Data S1**), they provided an opportunity to further evaluate the performance of our deep learning model. Our deep learning model that was trained by all six CDRs (see above) successfully predicted that 72 of the 81 (89%) antibodies as SARS-CoV-2 S antibodies (**Figure 6C and Table S5**). This result further demonstrates the possibility of predicting antibody specificity solely based on the primary sequence.

## DISCUSSION

Through a systematic survey of published information on SARS-CoV-2 antibodies, we identified many molecular features of public antibody responses to SARS-CoV-2. The large amount of published information has allowed us to explore distinct patterns of germline gene usages in antibodies that target different domains on the S protein (i.e. RBD, NTD, and S2). Notably, the types and nature of public antibody responses to different domains appear to be quite different. For example, convergence of CDR H3 sequences can be readily identified in the public antibody responses to RBD and S2. In contrast, the public antibody response to NTD seems to be largely independent of the CDR H3 sequence. Furthermore, an IGHD-dependent public antibody response was enriched against S2, but not RBD or NTD. Together, our study demonstrates the diversity of sequence features that can constitute a public antibody response against a single antigen.

The public antibody response to SARS-CoV-2 has also been examined by a recent data mining study that focused on identifying public clonotypes [55]. This previous study defined public clonotypes as antibodies with the same IGHV/IGHJ/IGK(L)V/IGK(L)V genes and high similarity of CDR H3 [55]. While multiple public clonotypes were identified using this stringent definition [55], the characterization of public antibody response is likely far from comprehensive. A public antibody response may not always involve a defined pair of IGHV/IGK(L)V genes, especially when either IGHV or IGK(L)V gene-encoded residues only make a minimal contribution to the paratope. In fact, a well-characterized public antibody response to the highly conserved stem region of influenza hemagglutinin has a paratope that is entirely attributed to the IGHV1-69 heavy chain [56-59]. IGHV3-30/IGHD1-26 antibodies to S2 in our study may represent a similar type of IGK(L)V-independent public antibody response, although it still needs to be confirmed by structural analysis. On the other extreme, RBD antibodies that are encoded by IGLV6-57 with a ^97^WLRG^100^ motif in the CDR H3 represent a public response that is largely independent of IGHV gene usage. Given the diverse types of public antibody responses to SARS-CoV-2 S, we need to acknowledge the limitation of using the conventional strict definition of public clonotype to study public antibody responses.

Public antibody response to different antigens can have very different sequence features. For example, IGHV6-1 and IGHD3-9 are signatures of public antibody response to influenza virus [24, 60-62], whereas IGHV3-23 is frequently used in antibodies to Dengue and Zika viruses [63]. In contrast, these germline genes are seldom used in the antibody response to SARS-CoV-2 as compared to the naïve baseline (**Figure 1B-C and Figure 3A**). Since the binding specificity of an antibody is determined by its structure, which in turn is determined by its amino acid sequence, the antigen specificity of an antibody can theoretically be identified based on its sequence. This study provides a proof-of-concept by training a deep learning model to distinguish between SARS-CoV-2 S antibodies and influenza HA antibodies, solely based on primary sequence information. Technological advancements, such as the development of single-cell high-throughput screen using the Berkeley Lights Beacon optofluidics device [64] and advances in paired B-cell receptor sequencing [65], have been accelerating the speed of antibody discovery and characterization. As more sequence information on antibodies to different antigens is accumulated, we may be able in the future to construct a generalized sequence-based model to accurately predict the antigen specificity of any antibody.

In summary, the amount of publicly available information on SARS-CoV-2 antibodies has provided invaluable biological insights that have not been readily obtained for other pathogens. One reason is that the COVID-19 pandemic has gathered scientists from many fields and around the globe to work intensively on SARS-CoV-2. The parallel efforts by many different research groups have enabled SARS-CoV-2 antibodies to be discovered in unprecedented speed and scale that have not been possible for other pathogens. We anticipate that knowledge of the molecular features of the antibody response to SARS-CoV-2 will keep accumulating as more antibodies are isolated and characterized. Ultimately, the extensive characterization of antibodies to the SARS-CoV-2 S protein may allow us to address some of the most fundamental questions about antigenicity and immunogenicity, as well as how the human immune repertoire has evolved to respond to specific classes of viral pathogens that have coexisted with humans for hundreds to thousands of years.

## MATERIALS AND METHODS

### Collection of antibody information

Information on the monoclonal antibodies is derived from the original papers (Supplementary Table 1). Sequences of each monoclonal antibody are from the original papers and/or NCBI GenBank database (www.ncbi.nlm.nih.gov/genbank) [52]. Putative germline genes were identified by IgBLAST [66]. Some studies isolated antibodies from multiple donors, but the donor identity for each antibody was not always clear. For example, some studies mixed B cells from multiple donors before isolating individual B cell clones. Since the donor identity cannot be distinguished among those antibodies, we considered them from the same donor with “_mix” as the suffix of the donor ID. In addition, the PBMCs of SARS-CoV survivors in three separate studies were all from NIH/VRC [12, 44, 67]. Since it is unclear If they are the same SARS-CoV survivor, the same donor ID “VRC_SARS1” was assigned to them to avoid overestimation of public antibody response. the neutralization activity of a given antibody was only measured at a single concentration, 50% neutralization activity or below was classified as non-neutralizing. We also downloaded the CoV-AbDab [2] in September 2021 to fill in any additional information. As of September 2021, there were 2,582 human SARS-CoV-2 antibodies in CoV-AbDab. Information in the finalized dataset was manually inspected by three different individuals. For antibodies that were shown to bind to S1 but not RBD, they were classified as NTD antibodies. Due to having identical nucleotide sequences, IGKV1D-39*01 was classified as IGKV1-39*01, IGHV1-68D*02 as IGHV1-68*02, IGHV1-69D*01 as IGHV1-69*19, IGHV3-23D*01 as IGHV3-23*01, and IGHV3-29*01 as IGHV3-30-42*01.

### Analysis of germline gene usages

Non-functional germline genes were ignored in our germline gene usage analysis. Except for the analysis presented in **Figure 1**, IGHV3-30-3 was classified as IGHV3-30 since they have identical amino-acid sequence in the framework regions, CDR H1, and CDR H2. To establish the baseline germline usage frequency, published antibody repertoire sequencing datasets from 26 healthy donors [31, 32] were downloaded from cAb-Rep [33]. Putative germline genes for each antibody sequence in these repertoire sequencing datasets from healthy donors were identified by were identified by IgBLAST [66].

### CDR H3 clustering analysis

Using a deterministic clustering approach, antibodies with CDR H3 sequences that had the same length and at least 80% amino-acid sequence identity were assigned to the same cluster. As a result, CDR H3 of every antibody in a cluster would have >20% difference in amino-acid sequence identity with that of every antibody in another cluster. A cluster would be discarded if all of its antibody members were from the same donor. The number of antibodies within a cluster was defined as the cluster size. Sequence logos were generated by Logomaker in Python [68]. For each cluster, epitope assignment was performed using the following scoring scheme. Briefly, there were three scoring categories, namely “RBD”, “NTD”, and “S2”.

- 1 point was added to category “RBD” for each antibody with an epitope label equals to “S:RBD” or “S:S1”.
- 1 point was added to category “NTD” for each antibody with an epitope label equals to “S:NTD”, “S:S1”, “S:non-RBD”, or “S:S1 non-RBD”.
- 1 point was added to category “S2” for each antibody with an epitope label equals to “S:S2”, “S:S2 Stem Helix”, “S:non-RBD”.

The category with >50% of the total points would be classified as the epitope for a given cluster. If no category had >50% of the total points, the epitope for the cluster would be classified as “unknown”.

### Identification of recurring somatic hypermutation (SHM)

In this study, a public clonotype was classified as antibodies from at least two donors that had the same IGHV/IGK(L)V genes and CDR H3s from the same CDR H3 cluster (see “CDR H3 clustering analysis” above). For each antibody, ANARCI was used to number the position of each residue according to Kabat numbering [69]. The amino-acid identity at each residue position of an antibody was then compared to that of the putative germline gene. CDR H3, CDR L3, and framework region 4 in both heavy and light chains were not included in this analysis. Insertions and deletions were also ignored in this analysis. SHM that occurred in at least two donors within a public clonotype was defined as a recurring SHM.

### Deep learning model for antigen identification

#### Model construction

The deep learning model consisted of two networks, namely multi-encoder (ME) and a stack of multi-layered perceptrons (MLP). The CDR amino-acid sequences were taken as input and passed to ME. Specifically, each CDR amino-acid sequence was described by a 21-letter alphabet vector 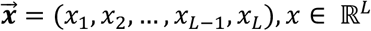, where L represented the length of sequence, and *x* represented the amino acid category. Each of the 20 canonical amino acids was one category, whereas all the ambiguous amino acids were grouped as the 21^st^ category. Before passing to ME, inputs were tokenized at the amino-acid level and processed by zero padding, so that the embedding layers represented the character-level tokens (i.e. amino acids) and the size of each input was the same. Subsequently, the inputs were mapped to the embedding vectors with additional dimension *d*. The sinusoidal positional encoding vectors were added to the embedding vectors to encode the relative position of tokens (i.e. amino acids) in the sequence. Each embedding vector, 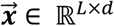, with size of *L* × *d*, was passed into transformer encoder layer by self-attention mechanism to learn the sequence feature [70]. All learned sequence features were then concatenated together and passed to multi-layered perceptron (MLP). Each MLP layer contained leaky rectified linear unit (ReLU) activations to avoid the vanishing gradient. Dropout layers were placed after each MLP block to avoid model overfitting [71]. The final output layer was followed by a sigmoid activation function to predict the probability of different classes. The prediction losses were calculated by binary cross-entropy loss.

#### Training detail

SARS-CoV-2 S antibodies and influenza HA antibodies with complete information for all six CDR sequences were identified. Sequences of each antibody were from the original papers (**Data S2**) or NCBI GenBank database (www.ncbi.nlm.nih.gov/genbank) [52]. If all six CDR sequences were the same between two or more antibodies, only one of these antibodies would be retained. After filtering duplicates, there were 4,736 antibodies to SARS-CoV-2 and 2,204 to influenza HA. The CDR sequences were identified by IgBLAST and PyIR [66, 72]. This dataset was randomly split into a training set (64%), a validation set (16%), and a test set (20%). The training set was used to train the deep learning model. The validation set was used to evaluate the model performance during training. The test set was used to evaluate the performance of the final model. There was no overlap of antibody sequences among the training set, validation set, and test set. The Adam algorithm was used to optimize the model. The following hyper-parameters were used for model training:

- CDR embedding size: 256
- The number of attention heads for self-attention on CDR feature learning: 4
- The number of encoder layer for CDR encoder: 4
- Size of stacking MLP layers: 512, 128, and 64
- Learning rate: 0.0001
- Batch size: 256

Using the same training set, validation set and test set, the model performance of using the following inputs was compared:

1. CDR H1 + H2
2. CDR L1 + L2
3. CDR H3
4. CDR L3
5. CDR H3 + L3
6. CDR H1 + H2 + H3
7. CDR L1 + L2 + L3
8. CDR H1 + H2 + H3 + L1 + L2 + L3

#### Performance Metrics

For evaluating model performance, S antibodies and HA antibodies were considered “positive” and “negative”, respectively. False positives (FP) and false negatives (FN) were samples that were misclassified by the model while true negatives (TN) and true positives (TP) were correctly classified one. The following metrics were computed to evaluate model performance:

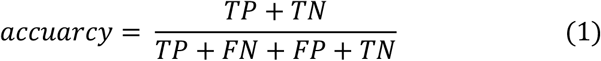

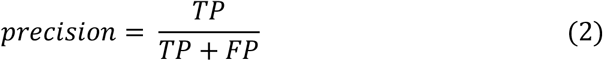

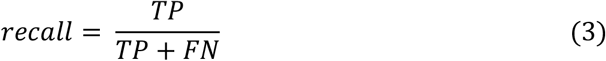

In addition, we also used the receiver operating characteristic (ROC) curve and precision-recall (PR) curve to measure the model’s ability to avoid misclassification [53, 54]. Area under the curves of ROC (i.e. ROC AUC) and PR (i.e. PR AUC) were computed using the “keras.metrics” module in TensorFlow [73].

## DATA AVAILABILITY

The assembled SARS-CoV-2 antibody dataset is in **Data S1**. The dataset for constructing and testing the deep learning model is in **Data S2**.

## CODE AVAILABILITY

Custom python scripts for all analyses have been deposited to https://github.com/nicwulab/SARS-CoV-2_Abs.

## ACKNOWLEDGEMENT

This work was supported by National Institutes of Health (NIH) R00 AI139445 (N.C.W.), DP2 AT011966 (N.C.W.), and Bill and Melinda Gates Foundation INV-004923 (I.A.W.). We thank Seth Zost and Huibin Lv for helpful discussion.

## AUTHOR CONTRIBUTIONS

All authors conceived and designed the study. Y.W, M.Y. and N.C.W. assembled the dataset and performed data analysis. J.P. provided technical expertise in deep learning, Y.W., M.Y. I.A.W, and N.C.W. wrote the paper and all authors reviewed and/or edited the paper.

