## Supplementary Figures for "A large-scale systematic survey of SARS-CoV-2 antibodies reveals recurring molecular features"

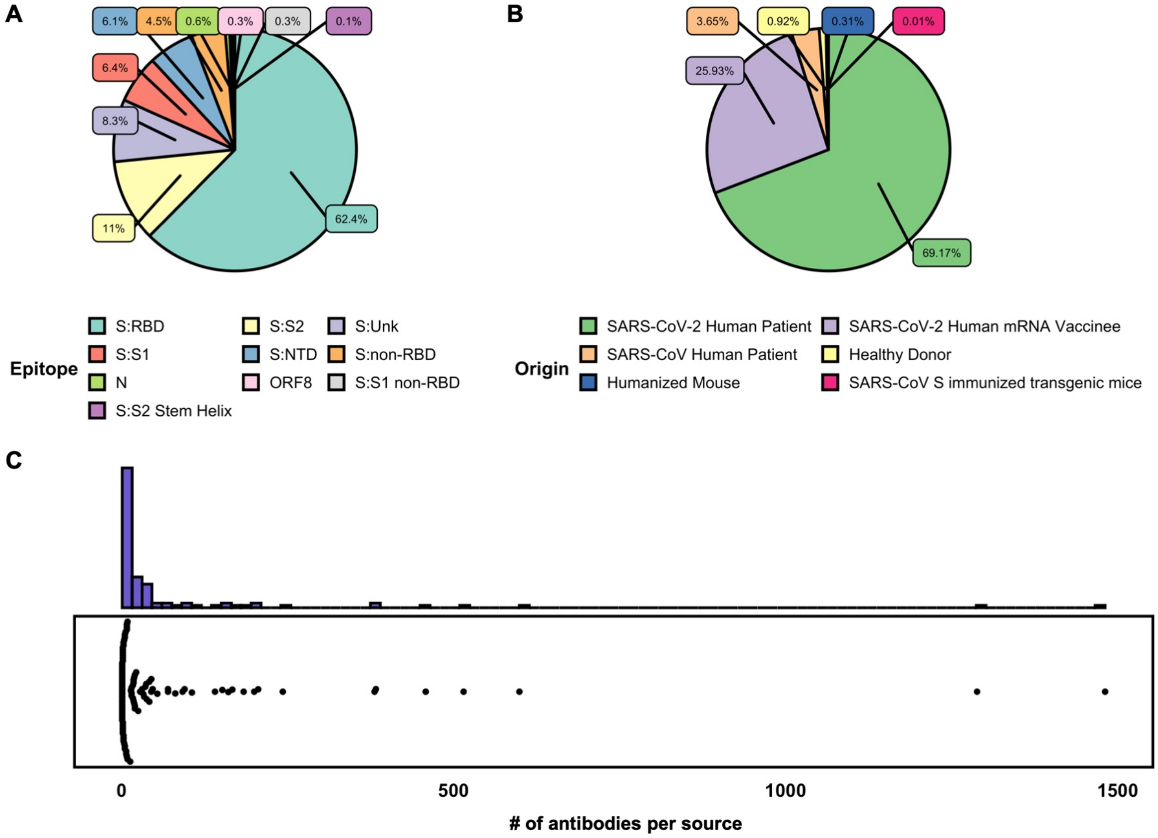


**Figure S1. Summary statistics of the antibody dataset. (A)** The percentages of antibodies in our antibody dataset that bind to different epitopes are shown. S:Unk represents spike protein with unknown binding domain. **(B)** The percentages of antibodies in our antibody dataset from different origins are shown. **(C)** Each data point in the bottom panel represents one source (i.e. one of the 88 research publications and 13 patents). The number of antibodies from each of the different sources is shown. The distribution of number of antibodies per source is also shown as a histogram in the upper panel.

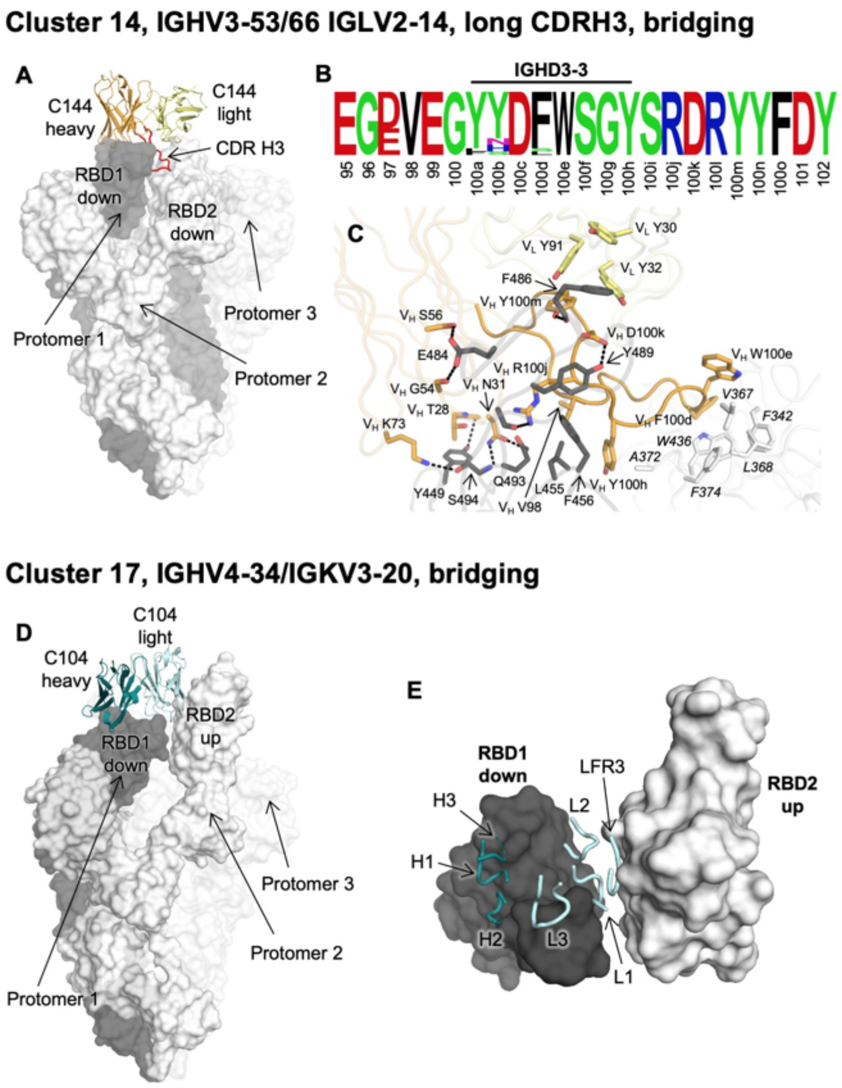


**Figure S2. CDR H3 clusters 14 and 17 represent antibodies that bridge two RBDs in the same spike trimer. (A-C)** Molecular feature of IGHV3-53/3-66 IGLV2-14 antibodies (cluster 14) that bridges two down RBDs simultaneously in a spike trimer. Cluster 14 contains 19 antibodies from 3 donors. **(A)** An overall view of SARS-CoV-2 spike in complex with an IGHV3-53/3-66 IGLV2-14 antibody. C144 (PDB 7K90) is used as a representative antibody [1]. Three protomers of the spike trimer are shown in charcoal, white, and transparent white, respectively. Heavy and light chains of C144 are shown in orange and yellow, respectively, with CDR H3 highlighted in red. The two RBDs bridged by the antibody are indicated in the figure. **(B)** Sequence logo of CDR H3 of IGHV3-53/3-66 IGLV2-14 antibodies, with residue positions labeled according to Kabat numbering. **(C)** Detailed interactions between C144 and the SARS-CoV-2 spike trimer. The color coding corresponds to that in panel A, where the RBD residues on protomer 1 is shown in charcoal and the RBD residues on protomer 2 are in white. Residue numbers of the RBD on protomer 2 are shown in italic. Hydrogen bonds and salt bridges are represented by black dashed lines. The RBD on protomer 1 is recognized by CDRs H1, H2, and H3 as well as a few light chain residues, where the RBD on protomer 2 is also targeted by CDR H3 of the same antibody molecule. CDR H3, which is encoded by IGHD3-3, contains $^{100d}$FW$^{100e}$ at tip that stacks extensively with aromatic residues in the adjacent RBD. **(D-E)** Molecular features of IGHV4-34/IGKV3-20 antibodies (cluster 17) bridging two RBDs (one up and one down) simultaneously in a spike trimer. Cluster 17 contains 13 antibodies from 4 donors. **(D)** An overall view of SARS-CoV-2 spike in complex with an IGHV4-34/IGKV3-20 antibody. C104 (PDB 7K8U) is used as a representative antibody [1]. Three protomers of the spike trimer are shown in charcoal, white, and transparent white, respectively. Heavy and light chains of C104 are shown in teal and pale cyan, respectively. The two RBDs bridged by the antibody are indicated in the figure. **(E)** A zoomed-in view of the interactions between the CDR loops of C104 and the two RBDs that displays one-up-one-down conformation. Atomic interactions are not shown here as side chains of paratope residues were truncated in the original structure due to the low resolution of the structure. CDR H3 was partially truncated in the original coordinates.

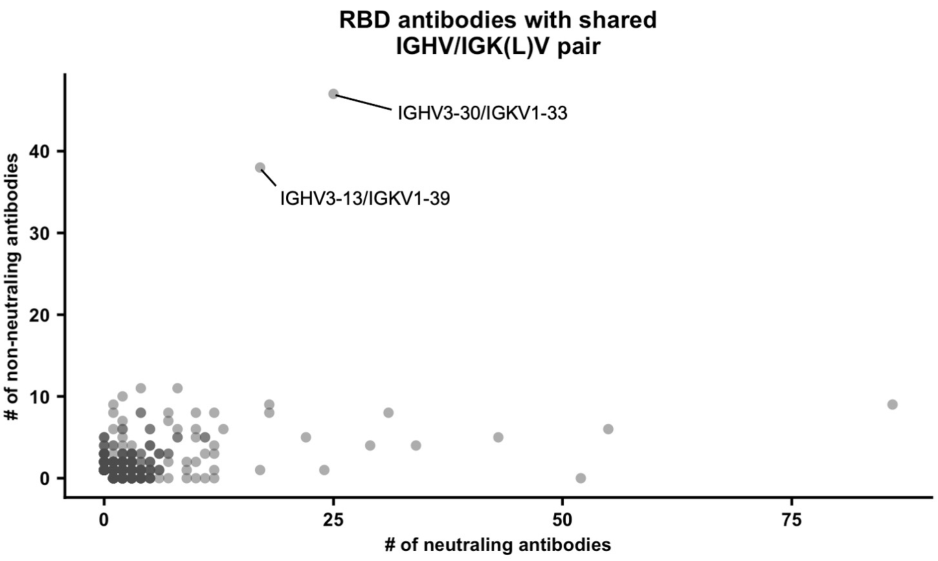


**Figure S3. Identification of IGHV/IGK(L)V pairs that are enriched in non-neutralizing RBD antibodies.** Each data point represents a defined IGHV/IGK(L)V pair. For each IGHV/IGK(L)V pair, the number of neutralizing RBD antibodies is plotted against the number of non-neutralizing RBD antibodies. Two IGHV/IGK(L)V pairs with a large number of non-neutralizing antibodies are labeled.

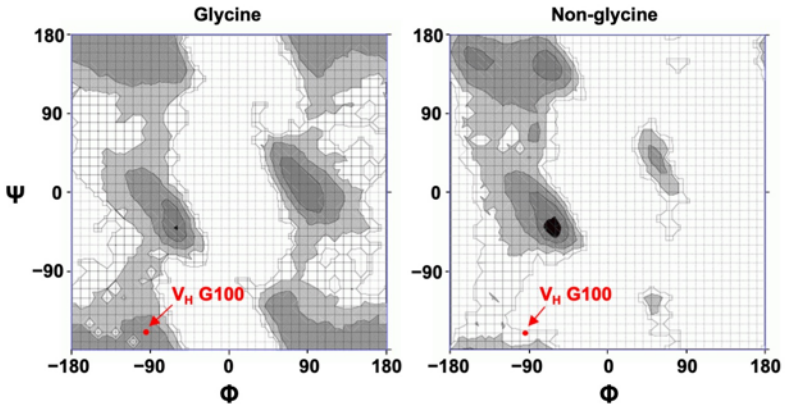


**Figure S4. Ramachandran plots for glycine and non-glycine amino acids.** $V_H$ G100 of antibody S2A4 (PDB 7JVA) [2], which is a representative antibody of cluster 7, is shown as red dots on the Ramachandran plots. Ramachandran plots were generated using the Ramachandran Plot Server (https://zlab.umassmed.edu/bu/rama/) [3]. Black and grey areas represent highly preferred conformations, whereas outlined white area represents allowed conformations.

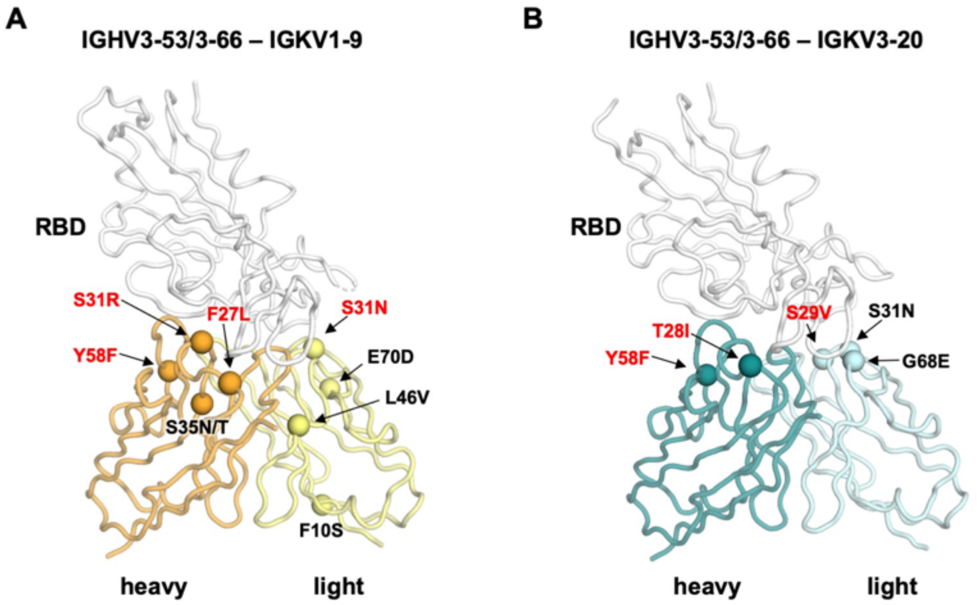


**Figure S5. Recurring somatic mutations in IGHV3-53/3-66 antibodies. (A)** Recurring somatic mutations in the public clonotype encoded by IGHV3-53/3-66 and IGKV1-9 are shown. CC12.1 (PDB 6XC3) is used as an example here [4]. **(B)** Recurring somatic mutations in the public clonotype encoded by IGHV3-53/3-66 and IGKV3-20 are shown. CC12.3 (PDB 6XC4) is used as an example here [4]. Paratope residues (defined as buried surface area upon binding > 0 Å$^2$ by RBD) are shown in red.
